# Exposure to 1800 MHz LTE electromagnetic fields under proinflammatory conditions decreases the response strength and increases the acoustic threshold of auditory cortical neurons

**DOI:** 10.1101/2022.01.13.476168

**Authors:** Samira Souffi, Julie Lameth, Quentin Gaucher, Délia Arnaud-Cormos, Philippe Lévêque, Jean-Marc Edeline, Michel Mallat

## Abstract

Increased needs in mobile phone communications have raised successive generations (G) of wireless technologies, which could differentially affect biological systems. We have assessed how a single 2h head-only exposure to a 4G long term evolution (LTE)-1800 MHz electromagnetic field (EMF) impacts on the microglial space coverage and the electrophysiological neuronal activity in the primary auditory cortex (ACx) in rats submitted to an acute neuroinflammation induced by lipopolysaccharide. The mean specific absorption rate in the ACx was 0.5 W/kg. Multiunit recording revealed that LTE-EMF triggered reduction in the response strength to pure tones and to natural vocalizations, together with an increase in acoustic threshold in the low and medium frequencies. Iba1 immunohistochemistry showed no change in the area covered by microglia cell bodies and processes. In healthy rats, the same LTE-exposure induced no change in response strength and acoustic threshold. Our data indicate that an acute neuroinflammation sensitize neuronal responses to LTE-EMF, which leads to an altered processing of acoustic stimuli in the ACx.

## Introduction

The electromagnetic environment of human populations has substantially evolved over the last three decades owing to the continuous expansion of wireless communications. Currently, more than two-thirds of the human population are considered as mobile phone (MP) users. The massive spreading of this technology has nourished concerns and debates about possible hazardous effect of pulsed electromagnetic fields (EMF) in the radiofrequency (RF) range, which are emitted by MP or base stations and encode the communications. This public health issue has stimulated a wealth of experimental research dedicated to the effects of RF absorption in biological tissues (SCENHIR, 2015). Parts of these investigations have looked for alterations of neuronal network activities and cognitive processes, considering the proximity of the brain to the source of RF under common use of MP. Many of the reported studies pertains to the influence of pulse-modulated signals used in the second generation (2G) global system for mobile communication (GSM), or in the wideband code-division multiple access (WCDMA) / 3^rd^ Generation Universal Mobile Telecommunication system (WCDMA /3G UMTS) (Curcio 2018; Kwon and Hamalainen 2011; Wallace and Selmaoui 2019; Zhang et al. 2017). Much less is known about the effects of RF signals used in the fourth generation (4G) mobile services, which relies on a fully digital Internet protocol – based technology called long term evolution (LTE) technology. The LTE mobile phone service was launched in 2011 and the worldwide number of LTE subscriptions was estimated to reach 5.43 billion in 2020 (Global mobile suppliers associations: //gsacom.com). In contrast with the GSM (2G) and WCDMA (3G) systems that are based on single-carrier modulation schemes, LTE uses orthogonal frequency division multiplexing (OFDM) as the basic signal format (Ghosh et al., 2010). Worldwide, LTE mobile services utilize a range of different frequency bands comprised between 450 MHz and 3700 MHz including the 900 and 1800 MHz bands also used in the GSM. The effect of an acute 30 min head exposure to a 2.573 GHz LTE signal on the global neuronal networks activity was recently explored in healthy human volunteers. Using resting state functional resonance magnetic imaging, it was observed that the LTE exposure induced alterations in spontaneous slow frequency fluctuations and in intraregional or interregional connectivities (Lv et al. 2014; Wei et al. 2019; Yang et al. 2021). Under similar exposure conditions, electroencephalographic analyses indicated reduced spectral powers and interhemispheric coherences in the alpha and beta bands (Yang et al. 2017). However, two other studies based on EEG analyses found that LTE head exposures had either no detectable effect (Nakatani-Enomoto et al. 2020) or resulted in reduced spectral power in the alpha band, with no alterations in cognitive functions assessed with the Stroop test (Vecsei et al. 2018). Significant discrepancies were also noted in the outcome of EEG or cognition studies dedicated to the effect of GSM or UMTS EMF exposures. They were thought to originate from variation in methodological designs and experimental parameters including signal types and modulations, exposure intensity and duration, or from heterogeneity among the human subjects with respect to age, anatomy or gender (Curcio 2018; Loughran et al. 2012; Zhang et al. 2017).

So far, animal studies have been seldom used to determine how exposure to LTE signals could affect brain functions. It was recently reported that whole body exposures of developing mice from late embryonic stages to weaning can cause altered locomotion and appetitive behaviors in adult life (Broom et al. 2019). Repeated exposures of adult rats were found to trigger oxidative stress and reduced the amplitude visual evoked potential obtained from the optic nerve (Ozdemir et al. 2021).

Besides analyses at multiple scales including cellular and molecular levels, rodent models can be used to investigate the effects of RF exposures during diseases, as illustrated by previous studies dedicated to the impact of GSM or WCDMA /3G UMTS EMF in the context of acute neuroinflammation, seizures, neurodegenerative diseases or glioma (Anane et al. 2003; Arendash et al. 2010; Carballo-Quintas et al. 2011; Jeong et al. 2015; Ouadah et al. 2018).

Lipopolysaccharide (LPS)-injected rodent is a classical preclinical model of an acute neuroinflammatory reaction associated to benign infectious diseases, which are caused by virus or bacteria and affect most of the human population each year. This inflammatory state is responsible for a reversible sickness and depressive behavior syndrome marked by fever, hypophagia and reduced social interactions (Dantzer et al. 2008). Residents CNS phagocytes such as microglia are key effector cells in this neuroinflammatory reaction. Rodent treatment with LPS triggers activation of microglia marked by remodeling of their shape and cell processes as well as deep changes in their transcriptome profile including upregulation of genes encoding proinflammatory cytokines or enzymes, which impact on the activity of neuronal networks (Hirbec et al. 2018; Hoogland et al. 2015; Kondo et al. 2011). Investigating the effects of a single head exposure to GSM-1800 MHz EMF in LPS-treated rats, we found that GSM signals trigger cell responses in the cerebral cortex affecting expression of genes, phosphorylation of glutamate receptors, neuronal evoked discharges and microglial cell morphology in the cerebral cortex. These effects were not detected in healthy rats submitted to a same GSM exposure indicating that the LPS-triggered neuroinflammatory state sensitize CNS cell responses to GSM signals (Lameth et al. 2020; Lameth et al. 2017; Occelli et al. 2018; Watilliaux et al. 2011). The capacity of RF exposure to impact on biological processes strongly depend on the specific absorption rate (SAR) expressed in W/kg, that measures the energy absorbed in the biological tissues. Focusing on the auditory cortex (ACx) of LPS-treated rats in which the local SAR averaged 1.55 W/kg, we observed that GSM-exposure resulted in an increased length or branching of microglia processes together with reduced neuronal responses evoked by pure and natural stimuli (Occelli et al., 2018).

In the current study, we aimed at examining whether a head-only exposure to LTE-1800 MHz signals could also alter microglial cell morphologies and neuronal activity in the ACx, reducing the power of the exposure by two-thirds. We show here that in the ACx of LPS-treated rats in which the average SAR value was 0.5 W/kg, LTE signals had no clear effect on microglial cell processes but, still, triggered significant reduction in sound-evoked cortical activity.

## Methods

### Subjects

Data were collected in the cerebral cortex of 31 adult male Wistar rats obtained from Janvier Laboratories at 55 days of age. Rats were housed in a humidity (50-55%) and temperature (22-24° C)-controlled facility on a 12 h/12 h light/dark cycle (light on at 7:30 A.M.) with free access to food and water. All experiments were conducted in accordance with the guidelines established by the European Communities Council Directive (2010/63/EU Council Directive Decree), which are similar to those described in the *Guidelines for the Use of Animals in Neuroscience Research of the Society of Neuroscience.* The protocol was approved by the ethical committee Paris-Sud and Centre (CEEA N°59, project 2014-25, national agreement 03729.02) using the procedures 32-2011 and 34-2012 validated by this committee.

The animals were habituated to the colony rooms for at least one week before LPS treatment and exposure (or sham-exposure) to LTE-EMF. Twenty-two rats were injected intraperitoneally (i.p.) with Ecoli LPS (250 μg/kg, serotype 0127:B8, SIGMA) diluted in sterile endotoxin-free isotonic saline 24h before LTE or sham-exposure (n= 11 per group).

### Exposure to LTE-1800MHz

Head-only exposure to LTE EMF was performed with an experimental setup previously used to assess the effect of GSM EMF (Lameth et al., 2017). LTE exposure was carried out 24h after LPS injections (11 animals) or without LPS treatment (5 animals). The animals were lightly anaesthetized with Ketamine/Xylazine (Ketamine 80 mg/kg, i.p; Xylazine 10 mg/kg, i.p) before exposure to prevent movements and ensure reproducible position of the animals’ head below the loop antenna emitting the LTE signal. Half of the rats from the same cages served as control (11 sham-exposed animals, among the 22 rats pretreated with LPS): they were placed below the loop antenna with the energy of the LTE signal set to zero. The weights of the exposed and sham-exposed animals were similar (p = 0.558, unpaired t-test, ns). All the anesthetized animals were placed on a metal-free heating pad to maintain their body temperature around 37°C throughout the experiment. As in previous experiments, the exposure duration was set at 2h. After exposure, the animals were placed on another heating pad in the surgery room. The same exposure procedure was also applied to 10 healthy rats (untreated with LPS), half of them from the same cages were sham-exposed (p = 0.694).

### EMF exposure system

The exposure system was analogous to the one described in previous studies (Leveque et al. 2004; Watilliaux et al. 2011), replacing the radiofrequency generator so as to generate LTE instead of GSM electromagnetic field. Briefly, a radiofrequency generator emitting a LTE −1800 MHz electromagnetic field (SMBV100A, 3.2 GHz, Rohde & Schwarz, Germany) was connected to a power amplifier (ZHL-4W-422+, Mini-Circuits, USA), a circulator (D3 1719-N, Sodhy, France), a bidirectional coupler (CD D 1824-2, −30 dB, Sodhy, France) and a four-ways power divider (DC D 0922-4N, Sodhy, France), allowing simultaneous exposure of four animals. A power meter (N1921A, Agilent, USA) connected to the bidirectional coupler allowed continuous measurements and monitoring of incident and reflected powers within the setup. Each output was connected to a loop antenna (Sama-Sistemi srl; Roma) enabling local exposure of the animal’s head. The loop antenna consisted of a printed circuit with two metallic lines engraved in a dielectric epoxy resin substrate (dielectric constant ε_r_ = 4.6). At one end, this device consisted of a 1 mm wide line forming a loop placed close to the animals’ head. As in previous studies (Lameth et al. 2017; Leveque et al. 2004), specific absorption rates (SARs) were determined numerically using a numerical rat model with the Finite Difference Time Domain (FDTD) method (Yee 1966; Kunz & Luebbers 1993; Taflove, 1995). They were also determined experimentally in homogeneous rat model using a Luxtron probe for measurement of rises in temperature. In this case, SARs, expressed in W/kg, were calculated using the following equation: SAR = C ΔT/Δt with C being the calorific capacity in J/(kg.K), ΔT, the temperature change in °K and Δt, the time in seconds. Numerically determined SAR values were compared with experimental SAR values obtained using homogenous models, especially in the equivalent rat brain area. The difference between the numerical SAR determinations and the experimentally detected SAR values was less than 30%.

Figure 1a shows the SAR distribution in the rat brain with a rat model, which matched that of the rats used in our study in terms of weight and size. The brain averaged SAR was 0.37 ± 0.23 W/kg (mean ± std). SAR values were the highest in cortical areas located directly below the loop antenna. The local SAR in the ACx (SAR_ACx_) was 0.50 ± 0.08 W/kg (mean ± std) (Fig. 1b). As the weight of the exposed rats was homogenous, differences in tissue thickness at the level of the head were negligible, so the actual SAR in ACx or other cortical regions was expected to be very similar from one exposed animal to another.

**Figure 1.**
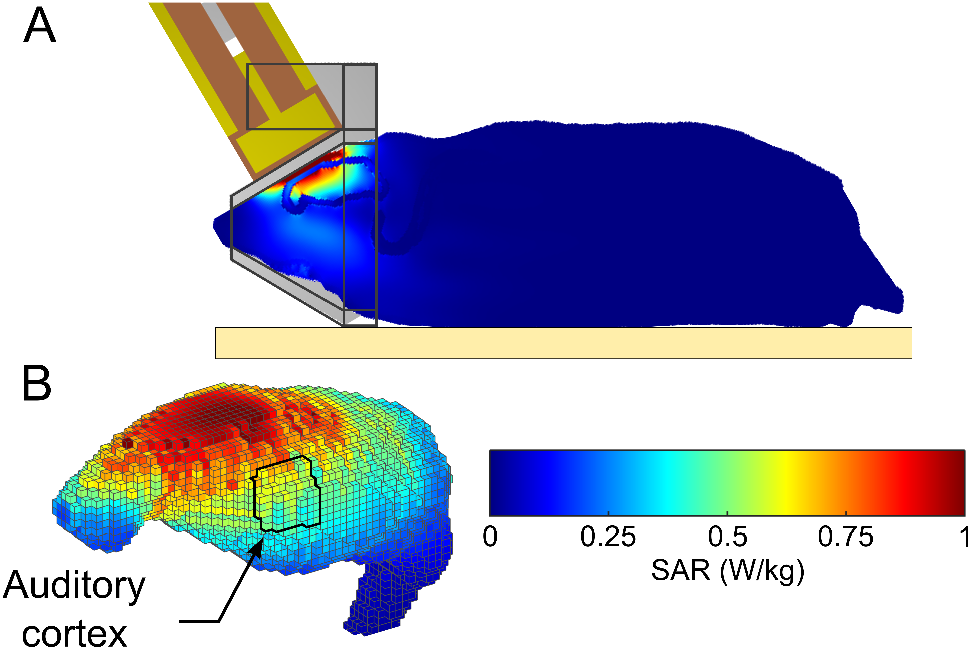
Dosimetric analysis of Specific Absorption Rates (SARs) in the rat brain during exposure to GSM 1800 MHz. The heterogeneous model of phantom rat and loop antenna described by Lévêque et al. (2004) was used to evaluate the local SAR in the brain with a 0.5 mm^3^ cubic mesh. (a) Global view of the rat phantom in the exposure setup with the loop antenna above its head and the metal-free heat pad (yellow) under the body. (b) Distribution of SAR values in the adult brain at 0.5 mm^3^ spatial resolution. The areas delimited by black contours in sagittal sections correspond to the primary auditory cortex in which microglia and neuronal activity were analyzed. The color-coded scale of SAR values applies to all numerical simulations shown in the figure.

### Surgical procedure for electrophysiological recordings

At the end of exposure, the animal was supplemented with additional doses of Ketamine (20 mg/kg, i.p.) and Xylazine (4 mg/kg, i.p.) until no reflex movement could be observed after pinching the hind paw. A local anesthetic (Xylocain 2%) was injected subcutaneously in the skin above the skull and on the temporal muscles. After placing the animal in a stereotaxic frame, a craniotomy was performed above the left temporal cortex. The opening was 9 mm wide starting at the intersection point between parietal and temporal bones and 5 mm height as in our previous studies (Cotillon and Edeline 2000; Cotillon-Williams and Edeline 2003; Manunta and Edeline 1997; Manunta and Edeline 1998; Manunta and Edeline 2004). The dura above the ACx was carefully removed under binocular control without damaging the blood vessels. At the end of the surgery, a pedestal in dental acrylic cement was built to allow atraumatic fixation of the animal’s head during the recording session. The stereotaxic frame supporting the animal was placed in a sound-attenuating chamber (IAC, model AC1).

### Recording procedures

Data were from multiunit recordings in the primary auditory cortex (area AI) of 20 rats including 10 animals pretreated with LPS. Extracellular recordings were from arrays of 16 tungsten electrodes (TDT, ø: 33 μm, <1 MΩ) composed of two rows of 8 electrodes separated by 1000 μm (350 μm between electrodes of the same row). A silver wire (ø: 300 μm), used as ground, was inserted between the temporal bone and the dura mater on the contralateral side. The estimated location of the primary ACx was 4-7 mm posterior to Bregma, 3 mm ventral to the superior suture of the temporal bone (corresponding area A1 in Paxinos & Watson 2005). The raw signal was amplified 10,000 times (TDT Medusa) then processed by a multichannel data acquisition system (RX5, TDT). The signal collected from each electrode was filtered (610-10000 Hz) to extract multi-unit activity (MUA). The trigger level was carefully set for each electrode (by a co-author blind to the exposed or sham-exposed status) to select the largest action potentials from the signal. On-line and off-line examination of the waveforms suggests that the MUA collected here was made of action potentials generated by 3 to 6 neurons at the vicinity of the electrode. At the beginning of each experiment, we set the position of the electrode array in such a way that the two rows of eight electrodes can sample neurons responding from low to high frequency when progressing in the rostro-caudal direction.

### Acoustic stimuli

Acoustic stimuli were generated in Matlab, transferred to a RP2.1-based sound delivery system (TDT) and sent to a Fostex speaker (FE87E). The speaker was placed at 2 cm from the rat’s right ear, a distance at which the speaker produced a flat spectrum (± 3 dB) between 140 Hz and 36 kHz. Calibration of the speaker was made using noise and pure tones recorded by a Bruel & Kjaer microphone 4133 coupled to a preamplifier B&K 2169 and a digital recorder Marantz PMD671. Spectro-temporal receptive fields (STRFs) were determined using 97 gamma-tone frequencies, covering eight (0.14-36 kHz) octaves presented at 75dB SPL at 4.15 Hz in a random order. The frequency response area (FRA) was determined using the same set of tones and presented from 75 dB to 5 dB SPL in random order at 2 Hz. Each frequency was presented eight times at each intensity.

Responses to natural stimuli were also evaluated. In previous studies (Occelli et al., 2018, 2019), we observed that, regardless of the neurons’ best frequency (BF), rat vocalizations rarely elicit robust responses in AI, whereas heterospecific ones (such as song birds or guinea pig vocalizations) often trigger robust and reliable responses across the entire tonotopic map. Therefore, we tested cortical responses to guinea pig’s vocalizations (a whistle call used in Gaucher et al. 2013a concatenated in a 1 sec stimuli, presented 25 times). This vocalization has its fundamental frequency between 2.5 and 5 kHz and harmonics up to 20 kHz (see Figure 2 in Gaucher et al., 2013a, and Figure 2 in Gaucher et al. 2013b).

**Figure 2.**
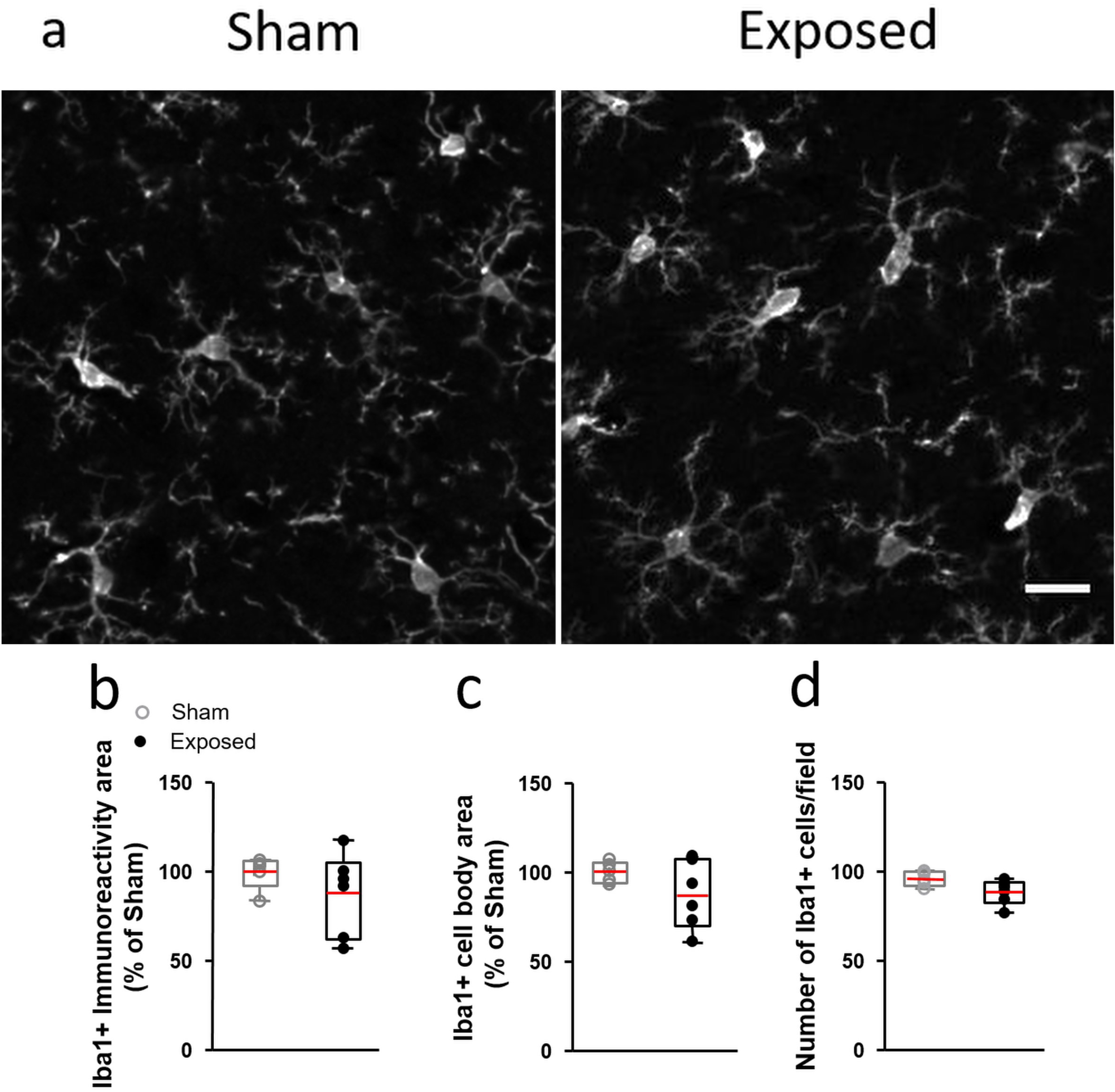
Microglia in the auditory cortex of LPS-injected rats after LTE or Sham exposure. (a) Representative stacked views of microglia stained with anti—Iba1 antibody in coronal sections of the auditory cortex from LPS—injected rats that were perfused 3 to 4h after Sham or LTE exposure (exposed). Scale bar: 20 μm. (b,c,d) Morphometric assessment of microglia 3 to 4h after sham (open dots) or LTE— exposure (exposed, dark dots). (b,c) Spatial coverage of the microglial marker Iba1(b) and area of Iba1-positive cell bodies (c). Data represents the area of anti-Iba1 staining normalized to mean values from Sham-exposed animals. (d) Counts of anti-Iba1-stained microglial cell bodies. Differences between Sham (n=5) and LTE (n=6) animals were not significant (p>0.05, unpaired t-test). The top and bottom of the box, and the upper and lower lines indicate 25^th^-75^th^ percentiles and 5^th^-95^th^ percentile respectively. The mean value is marked in red in the box.

Inserting an array of 16 electrodes in the cortical tissue systematically induces deformation of the cortex. We gave at least a 20-minute recovery period to allow the cortex to return to its initial shape, then the array was slowly lowered. Spectro-Temporal Receptive Fields (STRFs) were used to assess the quality of our recordings and to adjust the electrodes depth. The recording depth was 300-700 μm, which corresponds to layer III/IV and the upper part of layer V according to (Roger and Arnault 1989). When a clear tuning was obtained for at least 12 of the 16 electrodes, and that the stability of the recordings was satisfied for 15 minutes, the protocol started by presenting the acoustic stimuli in the following order: (1) Gamma-tones to determine the STRF at 75 dB SPL (5 min), (2) Gammatones at intensities ranging from 75 to 5 dB SPL (at 2 Hz) to determine the Frequency Responses Area (FRA, lasting 12min), (3) vocalizations (25 repetitions) at 75 dB SPL (2minutes). Presentation of the entire set of stimuli lasted 35 minutes, with periods of 2 min of spontaneous activity collected before and after these stimuli. This set of acoustic stimuli was used with the electrode array positioned at 3-5 locations per animal in the primary ACx.

### Data analysis

The STRFs derived from MUA were obtained by constructing post-stimulus time histograms (PSTHs) for each frequency with 1 ms time bins. All spikes falling in the averaging time window (starting at stimulus onset and lasting 100 ms) were counted. Thus, STRFs are matrices of 100 bins in abscissa (time) multiplied by 97 or 129 bins in ordinate (frequency). For visualization, all STRFs were smoothed with a uniform 5×5 bin window, but the actual values of response latency and bandwidth were quantified from raw data.

From the STRF obtained at 75dB SPL, the Best Frequency (BF) was defined as the frequency (in kHz) where the highest firing rate was recorded. At each intensity, peaks of significant response were automatically identified using the following procedure: A positive peak in the MUA-based STRF was defined as a contour of firing rate above the average level of the baseline activity (estimated from the ten first milliseconds of STRFs at all intensity levels) plus six times the standard deviation of the baseline (spontaneous) activity. For a given cortical recording, four measures were extracted from the peak of the STRF.

First, the “response strength” was the total number of spikes falling in the significant peaks of the STRFs. Second, the “response duration” was the time difference between the first and last spike of the significant peaks. Third, the “bandwidth” was defined as the sum of all peaks width in octaves. Forth, the response latency was measured as the time at which the firing rate was 6 times above spontaneous activity (i.e. the latency of the significant contour).

The response to the natural stimulus (guinea pig whistle) was quantified (i) by the firing rate (number of action potentials) emitted during the 345 ms of presentation of that stimulus and (ii) the trial-to-trial temporal reliability coefficient (CorrCoef) which quantifies the trial-to-trial reliability of the response over the 20 repetitions of the same stimulus. This index corresponds to the normalized covariance between each pair of spike trains recorded at presentation of this vocalization and was calculated as follows:

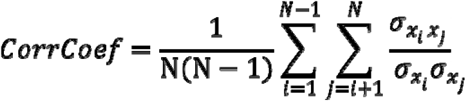

where *N* is the number of trials and *σ_xixj_* is the normalized covariance at zero lag between spike trains x_i_ and x_j_ where i and j are the trial numbers. Spike trains x_i_ and x_j_ were previously convolved with a 10-ms width Gaussian window, as in previous studies (Huetz et al., 2009; Gaucher et al., 2013a; Gaucher and Edeline, 2015; Aushana et al., 2018; Souffi et al., 2020, 2021). Based upon computer simulations, we have previously shown that this CorrCoef index is not a function of the neurons’ firing rate (Gaucher et al., 2013a).

### Immunohistochemistry

Eleven LPS-treated rats, submitted to identical LTE or sham exposure protocols (n = 6 or 5 per group), were used for immunohistochemistry in the cerebral cortex. Three to four hours after exposure (or sham exposure), adult rats were deeply anaesthetized with pentobarbital and perfused with 0.1 M phosphate buffered saline (PBS) followed by paraformaldehyde 4% in 0.1 M PBS, pH 7.4. Brains were post-fixated overnight in paraformaldehyde 4%, then cryoprotected by immersion in PBS containing 25% (w/v) sucrose and frozen in −70°C isopentane. Immunostaining was performed on coronal sections (20 μm thick) cut on a cryostat (Microm, Heidelberg, Germany) and mounted on gelatin-coated Superfrost glass slides (Menzel-glazer, Freiburg, Germany). Sections were blocked in PBS containing 0,1% Triton-X100 and 10% goat serum (30 min at room temperature) and then incubated overnight at 4°C with rabbit polyclonal anti-Iba1 (1:400, Wako Chemicals). Bound antibodies were detected by applying biotinylated goat anti-rabbit IgG (GE Healthcare, Velizy Villacoublay, France) for 2h at room temperature, followed by Alexa Fluor 488-conjugated streptavidin (Life technology, Saint Aubin, France) for 30 min at room temperature. Sections were counterstained with Hoechst 3342 dye (2μg/ml) and mounted in fluoromount G (Clinisciences). Control staining, performed by omitting primary antibodies, was negative.

### Image analysis

Quantitative image analyses were performed as described previously (Cheret et al. 2008). Images were captured using a Zeiss upright widefield microscope equipped with the apotome system. A 10× NA 0.50 Fluar objective (Zeiss microscope) was used to acquire images with equal exposure times. For each animal and each cortical region assessed, immunostained microglial cells were analyzed in microscopic fields (6 x 10^5^ μm^2^ area) acquired in 12 sections spanning areas of the auditory cortex and distributed over a cortical region extending in the rostro-caudal axis between 3.5 and 6.4 mm caudal to the bregma. Image analysis was performed blind relative to the experimental groups. The microglial cell bodies, confirmed by nuclei counterstaining, with Hoechst dye were counted manually. The area stained by anti-Iba1 and that of Iba1-stained cell bodies were determined following assignment of a threshold to eliminate background immunofluorescence, using Image J software.

### Statistical Analysis

Differences between groups of animals were analyzed using non-parametric Chisquare test or parametric unpaired Student t-test. Unpaired t-test was applied checking the assumption of equal variances and Gaussian distributions with F-test and Kolmogorov and Smirnov (K-S) test using Prism software.

## Results

### LTE exposure does not affect microglial cell morphology in the auditory cortex

Given previous demonstration that exposure to GSM-1800 MHz alters microglial cell morphology under proinflammatory conditions, we investigated this effect after exposure to LTE signals.

Adults rats were injected with LPS 24h before a 2h head-only Sham-exposure or an exposure to LTE-1800MHz with a mean SAR level of 0.5 W/kg in the ACx (Fig. 1). To determine whether LPS-activated microglia reacted to LTE EMF, we analyzed cortical sections stained with anti-Iba1 that selectively labels these cells (Kettenmann et al. 2011). As illustrated in Figure 2a microglial cells looked very similar in sections of ACx fixed 3 to 4h after sham or LTE exposure, showing a bushy like cell morphology marked by highly ramified tortuous processes. Consistent with an absence of morphological response, quantitative image analyses showed no significant differences in the total area of Iba1 immunoreactivity (unpaired t-test, p=0.308) or in the area (p=0,196) and the density (p=0.061) of Iba 1-stained cell bodies, when comparing LTE-exposed rats with Sham-exposed animals (Fig. 2b-d),

### LTE exposure decreases neuronal responses in LPS-treated animals

Table 1 summarizes the number of animals and multi-unit recordings obtained in the primary auditory cortex (A1) for the four groups of rats (Sham, Exposed, Sham-LPS, Exposed-LPS). In the following results, we have included all recordings exhibiting significant spectro-temporal receptive fields (STRFs), that is tone-evoked responses at least six standard deviations above spontaneous firing rate (see Table 1). Applying this criterion, we selected 266 recordings for the Sham group, 273 recordings for the Exposed group, 299 recordings for the Sham-LPS group and 295 recordings for the Exposed-LPS group.

In the following paragraphs, we will first describe the parameters extracted from the spectro-temporal receptive fields (i.e., the responses to pure tones) and from the responses to heterospecific vocalizations. Then, we will describe the quantifications of frequency response areas obtained for each group. To take into account the existence of “nested data” (see Aart et al., 2014) in our experimental design, all the statistical analyses were performed based on the number of positions of the electrode-array (last line in Table 1), but all the effects described below were also significant based upon the total number of multi-unit recordings collected in each group (third line in Table 1).

Figure 3a presents the distributions of best frequencies (BF, eliciting the largest responses at 75 dB SPL) for the cortical neurons obtained in the Sham and Exposed animals treated with LPS. The frequency range of the BFs extended from 1 kHz to 36 kHz in both groups. Statistical analysis showed that these distributions were similar (Chi-Square, p=0.278) indicating that comparisons between the two groups can be performed without sampling bias.

**Figure 3.**
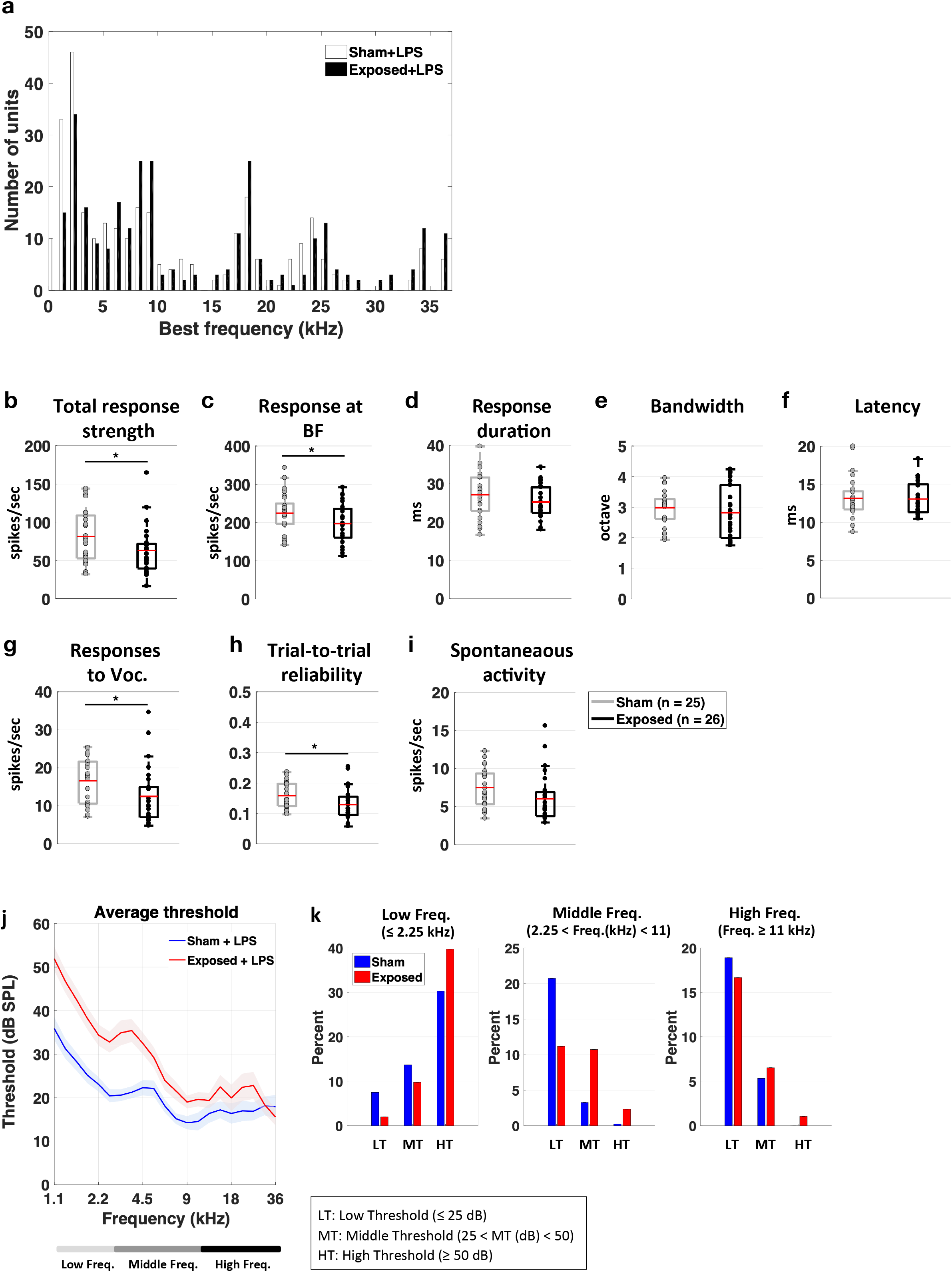
Effects of LTE exposure on parameters quantified from cortical responses in the LPS-treated animals. (a) BF distributions of the cortical neurons for the LPS-treated animals exposed to LTE (in black) and Sham-exposed to LTE (in white). The two distributions did not differ. (b-f) Effects of LTE exposure on the parameters quantifying the spectro-temporal receptive fields (STRF). The response strength was significantly decreased (* p< 0.05, unpaired t test) both on the entire STRF (total response strength) and at the best frequency (b-c). The response duration, the response bandwidth and the bandwidth were unchanged (d-f). Both the strength and the temporal reliability of the responses to vocalizations were decreased (g-h). Spontaneous activity was not significantly decreased (i). (* p<0.05, unpaired t-tests) (j-k) Effects of LTE exposure on the cortical thresholds. The average thresholds were significantly higher in LTE-exposed rats compared to Sham-exposed rats. This effect was more pronounced in the low and middle frequencies.

Figure 3b-f shows the distributions (means are represented by red lines) of the parameters derived from the STRFs for these animals. The effects induced by LTE exposure on the LPS-treated animals seem to point out a decrease in neuronal excitability. First, the total response strength and the response at the BF were significantly decreased compared to the Sham-LPS animals (Fig. 3b-c unpaired t tests, p=0.0017; and p=0.0445). Similarly, the responses to communication sounds were decreased both in terms of response strength and in terms of trial-to-trial reliability (Fig. 3g-h; unpaired t test, p=0.043). Spontaneous activity was decreased but this effect was not significant (Fig. 3i; p=0.0745). The response duration, the tuning bandwidth and the response latencies were not affected by LTE exposure in the LPS treated animals (Fig. 3d-f) suggesting that the frequency selectivity and the precision of onset responses were not impacted by LTE exposure in LPS-treated animals.

We next assessed whether the pure-tone cortical thresholds were altered by the LTE-exposure. Based on the Frequency Response Areas (FRAs) obtained from each recordings, we determined the auditory threshold at each frequency and averaged these thresholds in the two groups of animals. Figure 3j shows the mean (± sem) thresholds for the LPS treated rats from 1.1 to 36 kHz. Comparing the auditory thresholds of the Sham and Exposed groups revealed large increases in threshold in the Exposed animals compared to the Sham animals (Fig. 3j), an effect that was more prominent in the low and middle frequencies. More precisely, in low frequencies (<2.25 kHz), the proportions of A1 neurons with high thresholds increased and the proportion of low and mid-thresholds neurons decreased (Chi-Square = 43.85; p<0.0001; Fig. 3k, left panel). The same effect was also present in the middle frequencies (2.25<Freq(kHz)<11): there was higher proportion of cortical recordings with mid thresholds and a smaller proportion of neurons with low thresholds compared to the nonexposed group (Chi-Square=71.17; p<0.001; Fig. 3k, middle panel). There was also a significant difference in terms of threshold for the high frequency neurons (≥ 11 kHz, p=0.0059); the proportion of neurons with low threshold decreased and the proportion middle and high threshold increased (Chi-Square=10.853; p=0.04 Fig. 3k, right panel).

### LTE exposure did not change the response strength in healthy animals

Figure 4a presents the distributions of best frequencies (BF, eliciting the largest responses at 75 dB SPL) for the cortical neurons obtained in healthy animals for the Sham and Exposed group. Statistical analysis showed that these two distributions were similar (Chi-Square, p=0.157) indicating that comparisons between the two groups can be performed without sampling bias.

**Figure 4.**
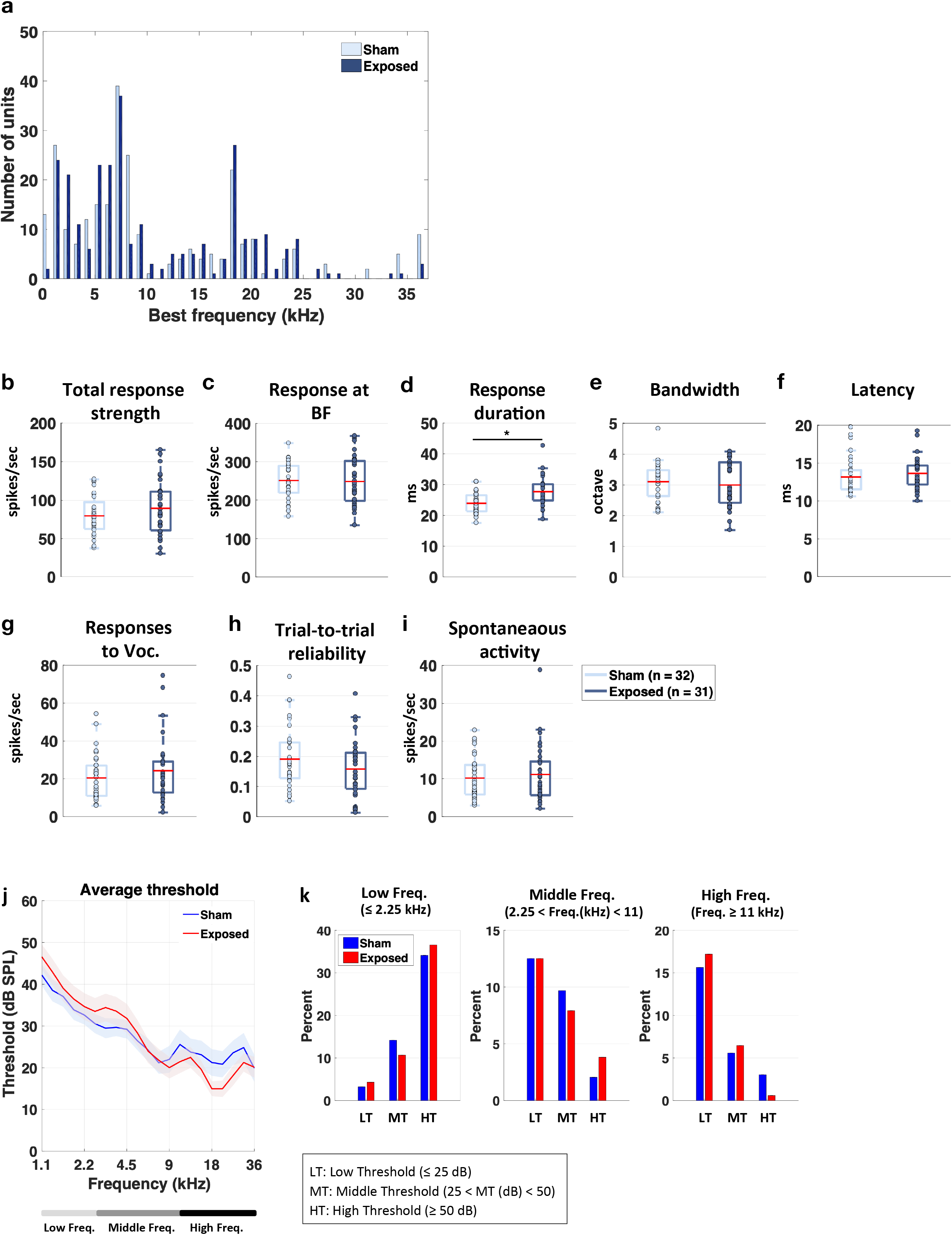
Effects of LTE exposure on parameters quantified from cortical responses in healthy animals. (a) BF distributions of the cortical neurons for healthy animals exposed to LTE (in dark blue) and Sham-exposed to LTE (in light blue). The two distributions did not differ. (b-f) Effects of LTE exposure on the parameters quantifying the spectro-temporal receptive fields (STRF). The response strength was not significantly changed both on the entire STRF and at the best frequency (b-c). The response duration was slightly increased (d) but, the response bandwidth and the bandwidth were unchanged (e-f). Both the strength and the temporal reliability of the responses to vocalizations were unchanged (g-h). Spontaneous activity was not significantly changed (i). (* p<0.05 unpaired t-tests) (j-k) Effects of LTE exposure on the cortical thresholds. On average, the threshold were not significantly changed in LTE-exposed rats compared to Sham-exposed rats, but the exposed animals had slightly lower thresholds in the high frequencies.

Figures 4b-f show box plots representing the distributions and mean values (red lines) of the parameters derived from the STRFs for the two groups. In healthy animals, LTE exposure by itself induced little change in the mean values of the STRF parameters. Compared to the Sham group (light blue boxes vs. dark blue boxes for the exposed group), the LTE exposure changed neither the total response strength nor the response at the BF (Fig. 4b-c; unpaired t-tests, p=0.2176, and p=0.8696 respectively). There was also no effect on the spectral bandwidth and the latency (respectively p=0.6764 and p=0.7129) but there was a significant increase in response duration (p=0.047). There was also no effect on the strength of responses to vocalizations (Fig 4g, p=0.4375), on the trial-to-trial reliability of these responses (Fig 4h, p=0.3412) and on the spontaneous activity (Fig 4i; p=0.3256).

Figure 4j shows the mean (± sem) thresholds for the healthy rats from 1.1 to 36 kHz. It did not reveal significant differences between the sham and exposed rats, except in the high frequencies (11-36 kHz) where the Exposed animals had slightly lower threshold (unpaired-t-test, p=0.0083). This effect reflects the fact that, in the exposed animals, there was slightly more neurons with low and middle thresholds (and less with high thresholds) in this frequency range (Chi-square=18.312, p=0.001; Fig. 4k).

To summarize, when performed on the healthy animals LTE-exposure has no effect on the response strength to pure tones and complex sounds such as vocalizations. In addition, in healthy animals the cortical auditory thresholds were similar between Exposed and Sham animals, whereas in the LPS-treated animals, the LTE exposure led to large increases in cortical thresholds especially in low and middle frequency ranges.

## Discussion

Our study reveals that in adult male rats undergoing acute neuroinflammation, an exposure to LTE-1800 MHz with a local SAR_ACx_ of 0.5 W/kg (see Methods) resulted in significant decreases in the strength of sound-evoked neuronal responses recorded in the primary ACx. These changes in neuronal activity occurred without any clear alteration in the extent of the spatial domain covered by microglial cell processes. This effect of LTE on the strength of cortical evoked responses was not observed in healthy rats. The differences in neuronal responsiveness can be attributed to biological effects of LTE signal rather than sampling bias, given the similarity of best frequency distributions among recorded units in LTE-exposed and sham-exposed animals (see Fig.3a). In addition, the absence of change in response latency and bandwidth of spectral tuning in LTE exposed rats indicates that, most likely, the recordings were sampled from the same cortical layers, which were localized in the primary ACx rather than in secondary area.

To our knowledge the effect of LTE signals on neuronal responses has not been reported previously. However, previous studies have documented the capacity of GSM-1800 MHz or 1800 MHz continuous wave (CW) to alter neuronal excitability, albeit with marked differences depending on the experimental approach. Recordings of snail ganglia shortly after an exposure to 1800 MHz CW at a SAR level of 8.2 W/Kg, showed reduction in the threshold for firing action potential and neuronal accommodation (Partsvania et al. 2013). On the other hand, spike and bursting activities in primary neuronal cultures derived from rat brain were reduced by exposures to GSM-1800 MHz or 1800 MHz CW applied for 15 minutes at a SAR of 4.6 W/kg. This suppressive effect was only partly reversible over a 30 min period after the end of the exposure. Full silencing of neurons was reached at a SAR of 9.2 W/kg. Dose-response analyses showed that GSM-1800 MHz was more efficient than 1800 MHz CW in suppressing burst activity indicating that neuronal responses depends on RF signal modulation (El Khoueiry et al. 2018).

In our setting, cortical evoked responses were collected *in vivo* three to six hours after the end of a 2h head-only exposure. In a previous study, we investigated the effect of GSM-1800 MHz at a SAR_ACx_ of 1.55 W/kg and found no significant effect on the sound-evoked cortical responses in healthy rats (Occelli et al., 2018). Here, the only significant effect induced in healthy rat by exposure to LTE-1800 at a SAR_ACx_ of 0.5 W/kg was a slight increase in response duration at pure tones presentation. This effect is difficult to explain as it was not accompanied by an increase in response strength indicating that this longer response duration occurred with the same total number of action potentials emitted by the cortical neurons. One explanation could be that the LTE exposure potentially reduced the activity of some inhibitory interneurons, as it is documented that in the primary ACx feedforward inhibitions control the duration of pyramidal cell responses triggered by excitatory thalamic inputs (Gaucher et al 2013, 2020; Ling et al. 2005; Tan et al. 2004; Wehr and Zador 2003).

In contrast, in rats submitted to LPS-triggered neuroinflammation, there was no effect of LTE exposure on the duration of sound-evoked neuronal firing, but marked effects were detected on the strength of evoked responses. Indeed, compared to neuronal responses recorded in LPS sham-exposed rats, neurons of LPS-treated rats exposed to LTE displayed a decrease in response strength, an effect observed both at presentation of pure tones or of natural vocalizations. The reduction in responses strength to pure tones occurred without narrowing the bandwidth of the spectral tuning at 75 dB, and as it occurred at all sound intensities, it induced an increase in acoustic threshold of cortical neurons in the low or medium sound frequencies.

The reduced strength of evoked responses indicates that the effects of LTE signal in LPS treated animals at a SAR_ACx_ of 0.5W/kg is similar to that of the GSM-1800 MHz applied at a three time higher SAR_ACx_ (1.55 W/kg; Ocelli et al., 2018). As for GSM signals, these results strongly suggest that head exposure to LTE-1800 MHz reduces neuronal excitability of ACx neurons in rats undergoing LPS-triggered neuroinflammation. In line with this assumption, we also observed a decrease in trial-to trial reliability of neuronal responses to vocalizations (Figure 3h) and a trend toward a reduction in spontaneous activity (Figure 3i). Obviously, it is quite difficult to determine *in vivo* if LTE signals reduce the neurons intrinsic excitability or reduce the synaptic inputs, which control neuronal responses in the ACx. Besides thalamic afferent projections that provide sound evoked inputs, it is now established that spectral inputs into a given cortical site also rely on cortico-cortical connections (Happel et al. 2010; Kaur et al. 2004; Metherate et al. 2005). However, the fact that we could not find significant difference in the bandwidth of cortical neurons between LTE- and sham exposed animals, potentially suggests that cortico-cortical connections did not play a major role in the LTE-induced decrease responses of ACx neurons. This also suggests that cortical inputs from other cortical regions exposed at higher SARs than those measured in ACx were unlikely to be responsible for the reported effects of LTE signal. In contrast, the LTE-induced increases in auditory threshold of ACx neurons raises the possibility that the thalamo-cortical transmission was reduced by the LTE exposure, because cortical thresholds strongly rely on thalamo-cortical synaptic inputs (Kaur et al, 2004; Happel et al, 2010).

Similar to the effects of GSM-1800 MHz, LTE-induced alterations of neuronal responses occurred in the context of LPS-triggered neuroinflammation, the signature of this being a microglial reaction (Hoogland et al., 2015). Current evidence indicate that microglia strongly impacts on the activity of neuronal networks both in the normal and in the pathological brain (Cantaut-Belarif et al. 2017; Kato et al. 2016; Merlini et al. 2021; Szalay et al. 2016). Their capacity to modulate neurotransmission relies not only on their production of compounds that potentialize, or limit, neurotransmission but also on the high motility of their cell processes. In the cerebral cortex, both enhancements and reductions in the activity of neuronal networks trigger rapid enlargement of microglia spatial domain due to the outgrowth of microglial cell processes (Liu et al. 2019; Umpierre et al. 2020). In particular, microglia protrusions are recruited in the vicinity of activated thalamo-cortical synapses and can suppress the activity of excitatory synapses through a mechanism involving microglia-mediated local generation of adenosine (Badimon et al. 2020).

In LPS-treated rats submitted to GSM-1800 MHz at a SAR_ACx_ of 1.55 W/kg, the reduced activity of ACx neurons occurred together with an outgrowth of microglia processes marked by a significant increase in Iba1-stained area in the ACx (Occelli et al., 2018). This observation suggested that microglial reshaping triggered by GSM exposure could actively contribute to the GSM-induced reduction in sound-evoked neuronal responses. Our current study argues against this hypothesis in the context of a LTE-head exposure with a SAR Acx limited to 0.5 W/kg, as we found no increase in the spatial domain covered by microglial processes. However, this does not rule out any effect of LTE signals on LPS activated-microglia, which could in turn affect neuronal activity. Further investigation is required to answer this question and to determine the mechanism by which acute neuroinflammation change the neuronal responsiveness to LTE signals.

To our knowledge the influence of LTE signals on auditory processing was not investigated previously. However, investigations carried out in humans or rodents found no consistent effect of GSM EMF on auditory processing at the cochlear or the brainstem level in healthy subjects (Arai et al. 2003; Aran et al. 2004; Bamiou et al. 2008; Galloni et al. 2005; Kwon et al. 2010; Oysu et al. 2005; Paglialonga et al. 2007; Parazzini et al. 2005; Sievert et al. 2005; Stefanics et al. 2007; Uloziene et al. 2005). Our previous studies (Occelli et al., 2018; Lameth et al., 2017) and the current one indicate that in the context of an acute inflammation, a head-only exposure to GSM-1800 MHz or LTE-1800MHz can result in functional alterations of neuronal responses in ACx, as indicated by the increase in auditory threshold of neuronal responses. These effects were observed 3 to 6h after the exposure and their persistence beyond 6h remain to be explored. However, we previously observed that in a cortical region receiving GSM signal at a SAR of 2.9 W/Kg, GSM-induced alterations in phosphorylation of excitatory glutamate receptors and in gene expressions faded out within 48h (Lameth et al., 2017). This suggests that effects described here of LTE exposure could be transient.

Our data can be put in perspective with respect to eligible SAR limits and estimates of actual SAR values reached in the cerebral cortex of mobile phone users. Current standards applied to protect the general public set the SAR limit to 2 W/kg for a local head or torso exposure to RF ranging between 100 kHz and 6 GHz RF (ICNIRP, 2020).

Dosimetric simulations have been performed using different human head models to determine RF power absorption in the head in general, or in the different tissues of the head during a mobile phone communication. Besides the diversity in the human head models, the simulations emphasize significant differences or uncertainties in the estimations of the energy absorbed in the brain according to anatomical or histological parameters such as the external or internal shape of the skull, the thickness or the water content of the different head tissues, which can strongly vary according to age, gender or across individuals (Christ et al. 2010; Fernandez-Rodriguez, 2015; Lee et al. 2019; Wiart et al. 2008). Furthermore, cell phone features such as the internal location of the antenna and cell phone positions relative to the user’s head, strongly impact on the levels and distribution of SAR values in the cerebral cortex (Belrhiti et al., 2017; Cardis et al. 2008; Ghanmi et al. 2014). However, considering the reported SAR distribution over the human cerebral cortex, which were established with from phone models emitting RF in the range of 1800 MHz (Cardis, 2008; Ghanmi, 2014; Lee, 2019), it appears that SAR levels reached in the human auditory cortex remained less than half those applied in our study (SAR_ACx_ 0.5 W/kg). Therefore, our data do not challenge the current limitations in SAR values applied to the general public.

In conclusion, our study reveals that a single head-only exposure to LTE-1800 MHz can interfere with the neuronal responses of cortical neurons to sensory stimuli. In line with previous characterizations of the effect of GSM-signal, our results show that the impact of LTE signal on neuronal activity varies according to the health state. Acute neuroinflammation sensitize neuronal responses to LTE-1800 MHz, resulting in altered cortical processing of auditory stimuli.

## Acknowledgements

We thank Icm.Quant core facility for assistance with image analysis. This research was supported by the French National Research Program for Environmental and Occupational health of ANSES (2018/2 RF/16) and funding from the program “Investissements d’avenir” ANR-10-IAIHU-06.

## Declarations

### Availabililty of Data and material

Data used in this study are available on reasonable request

### Code availability

Not applicable

### Ethical Approval

All procedures and experiments were conducted in accordance with the guidelines established by the European Communities Council Directive (2010/63/EU Council Directive Decree), which are similar to those described in the *Guidelines for the Use of Animals in Neuroscience Research of the Society of Neuroscience.* The protocols were approved by the institutional ethic committee Paris-Sud and Centre (CEEA N°59, project 2014-25, national agreement 03729.02) using the procedures 32-2011 and 34-2012 validated by this committee. We further attest that all efforts were made to minimize the number of animal used and their suffering.

### Consent to participate

Not applicable

### Consent to publication

Not applicable

